## Supplementary Figures and Tables for "Joint representation of molecular networks from multiple species improves gene classification"

#### Supplemental Material

#### Multi-species networks

**Table SM1** shows the properties of networks from multiple species considered in this study, obtained from BioGRID (Stark *et al.*, 2006) (v3.5.175) and IMP (Wong *et al.*, 2015) (v2). Note that zebrafish was included only in some analyses in this work due to its small network size in BioGRID and limited number of known gene annotations to biological processes and phenotypes.

**Table SM1: Network Coverage and Density.** Genes: Number of genes in the network, Edges: Total number of edges in the network, Density: Ratio of number of edges / total possible edges

| Species | BioGRID Number of Genes | BioGRID Number of Edges | BioGRID Density | IMP Number of Genes | IMP Number of Edges | IMP Density |
|---|---|---|---|---|---|---|
| human | 17,692 | 338,669 | 0.0022 | 28,443 | 77,361,677 | 0.19 |
| mouse | 7,296 | 20,673 | 0.0008 | 36,242 | 250,547,482 | 0.38 |
| zebrafish | 208 | 216 | 0.01 | 22,646 | 64,559,106 | 0.25 |
| fly | 9,191 | 60,280 | 0.0014 | 20,397 | 112,978,222 | 0.54 |
| worm | 6,210 | 23,358 | 0.0012 | 21,030 | 55,513,529 | 0.25 |
| yeast | 6,144 | 537,204 | 0.0285 | 6,291 | 4,086,174 | 0.21 |

The first step in GenePlexusZoo is to connect gene networks from different species by adding an edge between pairs of genes if they belonged to the same EggNOG orthologous group (Huerta-Cepas *et al.*, 2019). For each species, **Figure SM1** shows the number and fraction of genes in that species network that is part of a cross-species edge when combined with a network from another species.

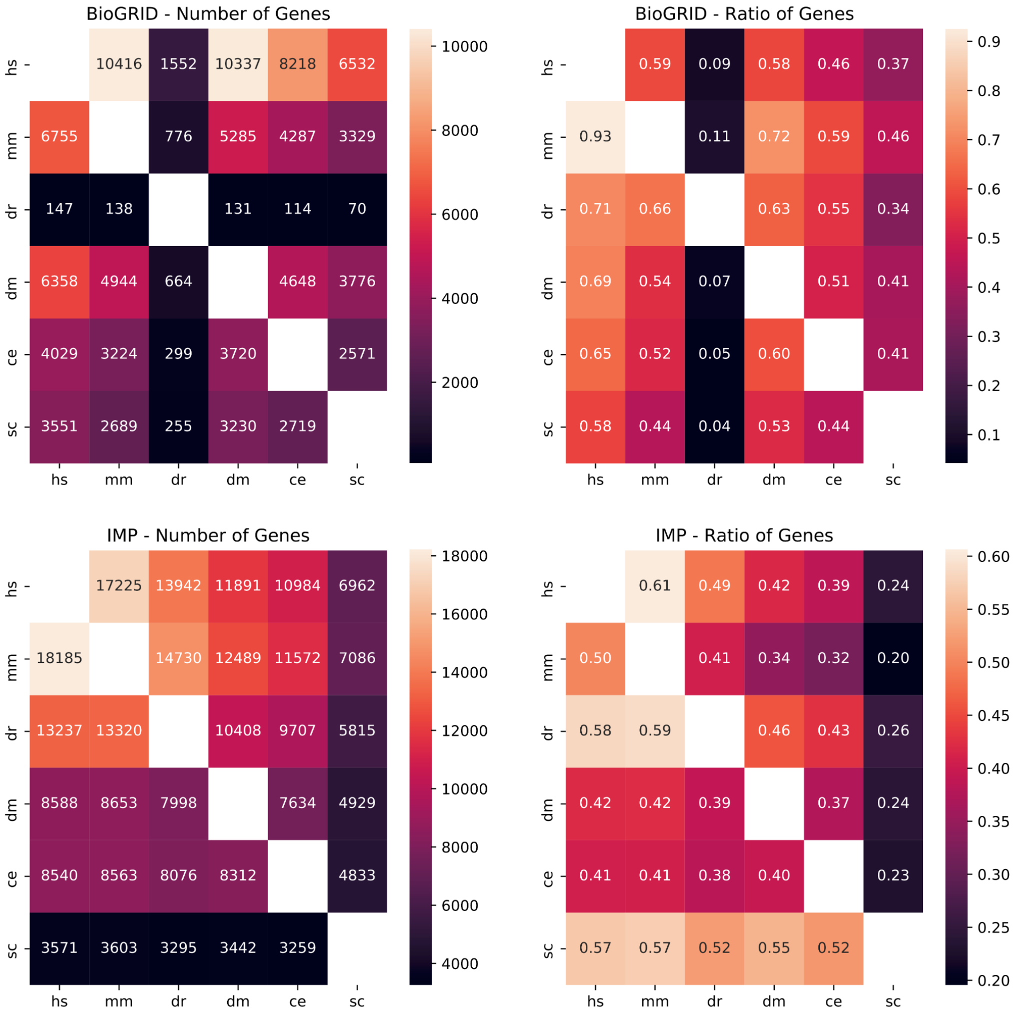


**Figure SM1: Number and ratio of cross-species network connections.** A row in each heat map in the left column shows the total number of genes in one species (row) present in a network of species combined with the network of another species (column). The corresponding entries in each heat map in the right column show the fraction of genes with a cross-species edge (when combined with a network from one other species). The top and bottom rows correspond to networks from BioGRID and IMP, respectively.

These cross-species gene links were weighted in two different ways (**Fig. SM2**; which were then compared). In the 'uniform edge weighting strategy', the edge weight was set to the same value for all cross-species edges. We devised a second 'degree weighting strategy' that weights edges differently based on the degrees of the incident gene nodes in their respective within-species networks. Specifically, for a pair of genes $u$ and $v$ in two different species, the cross-species edge weight, $w_c$, is given by:

$$w_c(u, v) = \frac{\left(C * W_{within}\right)}{N_{orthos}}, \qquad \text{(eqn. SM1)}$$

Here, $W_{within}$ is the degree of the source node *u* when only considering within-species edges, $N_{orthos}$ is the total number of ortholog connections for *u*, and *C* is a scaling parameter. The parameter *C* influences the *node2vec* in how likely it is to generate walks that, after landing on node *u,* will next traverse to a node in the same species or to a node in a different species.

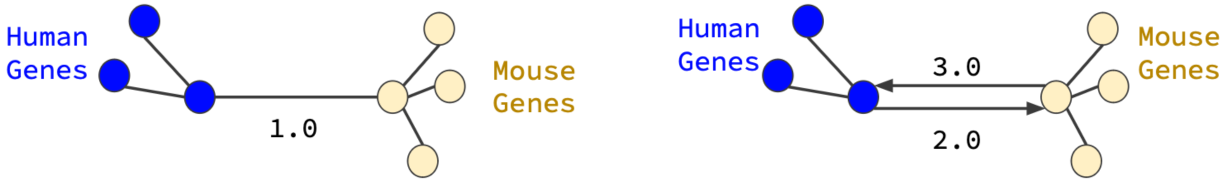


**Figure SM2: Edge weighting strategies for connecting genes across species.** The uniform edge weighting strategy sets all cross species edges to the same value (left). The degree weighting strategy (shown with C = 1) creates directed edges by considering both the within-species degree of the source node and the total number of cross-species edges the source node is incident on (right).

### Creating multi-species network representations

After connecting networks from multiple species, we derived representations (to serve as features for ML) using four different methods:

1. **AdjMat**: Adjacency matrix representation of the networks where the features directly use the edge weights for all the connections. That is, each gene's feature vector is its corresponding row in the adjacency matrix.
2. **RWR**: The influence matrix representation generated with a random-walk-with-restart kernel (Leiserson *et al.*, 2015).
    - The influence matrix $F = \beta[I - (1 - \beta)W_D]^{-1}$, where, β is the restart parameter, $I$ is the identity matrix, and $W_D$ is the degree-weighted adjacency matrix given by $W_D = AD^{-1}$, where $D \in R^{|V|x|V|}$ is a diagonal matrix of node degrees. We used either the influence matrix directly or its transpose.
3. **SVD**: Embedding of the adjacency matrix using SVD with or without column-wise standardization of the adjacency matrix before SVD.
4. **N2V**: Embedding of the adjacency matrix using *node2vec* (Grover and Leskovec, 2016).

### Properties of gene set collections

This section reports the properties of the various gene set collections. The following information is contained in these tables (* denotes wildcard expression):

- **Task**: Name of the gene set collection; Gene Ontology (GO) (Ashburner *et al.*, 2000), Monarch (MO) (Mungall *et al.*, 2017), and DisGeNet (DGN) (Piñero *et al.*, 2017; Schriml *et al.*, 2019).
- **Species**: Species for which the annotations of for; hs: human, mm: mouse, dr: zebrafish, dm: fly, ce: worm, sc: yeast

- **Network**: Network from which features are derived. Bio : BioGRID
- **NumSets**: Total number of sets that passed the threshold
- **All***: Considering all genes annotated to the set
- **Trn***: Considering only annotated genes in the training set
- **Tst***: Considering only genes in the testing set
- **Min**: The number of genes annotated to the gene set with the least amount of annotations
- **Max**: The number of genes annotated to the gene set with the greatest amount of annotations
- **Mean**: Average number of genes annotated to sets in the collection
- **Median**: Median value of the number of genes annotated to sets in the collection

**Table SM2: Information on the non-redundant sets.**

| Task | Species | Network | NumSets | All-Min | All-Max | All-Mean | All-Median | Trn-Min | Trn-Max | Trn-Mean | Trn-Median | Tst-Min | Tst-Max | Tst-Mean | Tst-Median |
|---|---|---|---|---|---|---|---|---|---|---|---|---|---|---|---|
| GO | ce | Bio | 58 | 24 | 148 | 75.9 | 70.5 | 12 | 122 | 59.7 | 57.5 | 10 | 44 | 16.2 | 14.5 |
| GO | dm | Bio | 133 | 23 | 188 | 88.7 | 87.0 | 12 | 174 | 71.3 | 71.0 | 10 | 77 | 17.5 | 15.0 |
| GO | hs | Bio | 201 | 21 | 100 | 62.1 | 64.0 | 10 | 87 | 45.5 | 46.0 | 10 | 51 | 16.6 | 14.0 |
| GO | mm | Bio | 76 | 21 | 78 | 45.7 | 44.5 | 10 | 64 | 31.8 | 31.5 | 10 | 29 | 13.9 | 12.0 |
| GO | sc | Bio | 113 | 20 | 198 | 84.0 | 75.0 | 10 | 182 | 60.9 | 52.0 | 10 | 83 | 23.1 | 17.0 |
| MO | ce | Bio | 52 | 33 | 147 | 87.2 | 86.5 | 23 | 123 | 70.7 | 71.0 | 10 | 41 | 16.5 | 15.5 |
| MO | dm | Bio | 3 | 149 | 185 | 169.7 | 175.0 | 139 | 173 | 158.0 | 162.0 | 10 | 13 | 11.7 | 12.0 |
| MO | hs | Bio | 208 | 20 | 99 | 60.0 | 57.5 | 10 | 87 | 45.8 | 43.0 | 10 | 30 | 14.1 | 13.0 |
| MO | mm | Bio | 82 | 22 | 70 | 41.1 | 40.5 | 11 | 59 | 28.6 | 27.0 | 10 | 20 | 12.5 | 12.0 |
| MO | sc | Bio | 50 | 30 | 196 | 108.4 | 101.0 | 11 | 172 | 87.1 | 78.0 | 10 | 97 | 21.3 | 14.5 |
| DGN | hs | Bio | 140 | 26 | 189 | 105.5 | 106.5 | 10 | 171 | 85.4 | 83.0 | 10 | 75 | 20.1 | 16.0 |
| GO | ce | IMP | 100 | 20 | 198 | 87.3 | 79.0 | 10 | 183 | 68.2 | 59.5 | 10 | 69 | 19.1 | 15.5 |
| GO | dm | IMP | 107 | 28 | 188 | 98.1 | 94.0 | 15 | 164 | 80.9 | 77.0 | 10 | 69 | 17.3 | 15.0 |
| GO | dr | IMP | 112 | 20 | 189 | 87.8 | 75.5 | 10 | 153 | 66.8 | 57.5 | 10 | 76 | 21.0 | 17.5 |
| GO | hs | IMP | 210 | 23 | 100 | 64.4 | 65.0 | 10 | 90 | 47.2 | 47.0 | 10 | 62 | 17.2 | 14.0 |
| GO | mm | IMP | 220 | 23 | 99 | 64.9 | 68.0 | 10 | 87 | 49.2 | 51.5 | 10 | 52 | 15.7 | 13.0 |
| GO | sc | IMP | 109 | 20 | 198 | 85.1 | 76.0 | 10 | 183 | 62.7 | 56.0 | 10 | 79 | 22.4 | 17.0 |
| MO | ce | IMP | 88 | 22 | 189 | 95.4 | 88.5 | 11 | 166 | 77.7 | 67.5 | 10 | 40 | 17.7 | 15.0 |
| MO | dm | IMP | 25 | 33 | 194 | 106.8 | 108.0 | 15 | 168 | 81.9 | 73.0 | 10 | 89 | 24.9 | 17.0 |
| MO | dr | IMP | 119 | 23 | 187 | 67.9 | 56.0 | 10 | 166 | 48.1 | 39.0 | 10 | 76 | 19.8 | 15.0 |
| MO | hs | IMP | 160 | 20 | 96 | 56.9 | 55.0 | 10 | 85 | 41.2 | 39.0 | 10 | 33 | 15.7 | 14.0 |
| MO | mm | IMP | 176 | 22 | 100 | 64.4 | 65.0 | 10 | 89 | 49.4 | 50.0 | 10 | 42 | 15.0 | 13.0 |
| MO | sc | IMP | 50 | 30 | 196 | 108.7 | 101.0 | 11 | 172 | 87.3 | 79.0 | 10 | 97 | 21.4 | 14.5 |
| DGN | hs | IMP | 143 | 22 | 199 | 107.4 | 106.0 | 10 | 179 | 87.5 | 82.0 | 10 | 74 | 19.9 | 16.0 |

**Table SM3. Information on the matched non-redundant sets.**

| Task | Species | Features | Network | NumSets | All-Min | All-Max | All-Mean | All-Median | Trn-Min | Trn-Max | Trn-Mean | Trn-Median | Tst-Min | Tst-Max | Tst-Mean |
|---|---|---|---|---|---|---|---|---|---|---|---|---|---|---|---|
| GO | ce | ce_hs | Bio | 16 | 21 | 126 | 70.6 | 69.5 | 9 | 100 | 56.6 | 56.0 | 10 | 26 | 14.0 |
| GO | dm | dm_hs | Bio | 66 | 17 | 188 | 78.0 | 65.0 | 7 | 166 | 61.5 | 47.5 | 10 | 58 | 16.4 |
| GO | hs | ce_hs | Bio | 16 | 29 | 192 | 113.9 | 120.5 | 16 | 164 | 89.8 | 95.0 | 10 | 47 | 24.1 |
| GO | hs | dm_hs | Bio | 66 | 38 | 190 | 124.5 | 131.0 | 12 | 168 | 97.1 | 99.0 | 10 | 82 | 27.4 |
| GO | hs | hs_mm | Bio | 80 | 19 | 195 | 129.4 | 137.5 | 8 | 176 | 106.3 | 113.5 | 10 | 48 | 23.1 |
| GO | hs | hs_sc | Bio | 92 | 16 | 190 | 99.8 | 95.0 | 6 | 170 | 72.1 | 67.5 | 10 | 83 | 27.7 |
| GO | mm | hs_mm | Bio | 80 | 18 | 141 | 66.0 | 64.5 | 6 | 116 | 51.6 | 47.0 | 10 | 35 | 14.3 |
| GO | sc | hs_sc | Bio | 92 | 15 | 170 | 64.5 | 52.5 | 5 | 151 | 43.6 | 34.5 | 10 | 74 | 21.0 |
| GO | ce | ce_hs | IMP | 30 | 20 | 163 | 68.6 | 57.0 | 9 | 134 | 53.1 | 41.5 | 10 | 56 | 15.6 |
| GO | dm | dm_hs | IMP | 53 | 20 | 197 | 84.1 | 67.0 | 7 | 177 | 67.7 | 51.0 | 10 | 69 | 16.4 |
| GO | dr | dr_hs | IMP | 24 | 20 | 187 | 84.4 | 68.5 | 8 | 142 | 62.4 | 46.5 | 10 | 45 | 22.0 |
| GO | hs | ce_hs | IMP | 30 | 32 | 199 | 130.9 | 150.5 | 15 | 168 | 98.7 | 109.5 | 12 | 89 | 32.2 |
| GO | hs | dm_hs | IMP | 53 | 20 | 196 | 120.4 | 125.0 | 5 | 163 | 90.1 | 86.0 | 10 | 89 | 30.3 |
| GO | hs | dr_hs | IMP | 24 | 37 | 197 | 117.3 | 107.0 | 13 | 174 | 93.8 | 79.5 | 10 | 71 | 23.5 |
| GO | hs | hs_mm | IMP | 198 | 20 | 198 | 122.9 | 129.0 | 5 | 179 | 97.0 | 98.0 | 10 | 87 | 25.8 |
| GO | hs | hs_sc | IMP | 87 | 16 | 196 | 105.3 | 102.0 | 6 | 175 | 76.0 | 76.0 | 10 | 89 | 29.2 |
| GO | mm | hs_mm | IMP | 198 | 17 | 200 | 88.4 | 81.0 | 5 | 180 | 70.3 | 62.0 | 10 | 62 | 18.1 |
| GO | sc | hs_sc | IMP | 87 | 17 | 170 | 65.4 | 50.0 | 7 | 154 | 44.7 | 35.0 | 10 | 73 | 20.8 |

**Table SM4. Information on the full gene set collections.**

| Task | Species | Network | NumSets | All-Min | All-Max | All-Mean | All-Median |
|---|---|---|---|---|---|---|---|
| GO | ce | Bio | 967 | 10 | 165 | 33.8 | 22 |
| GO | dm | Bio | 2002 | 10 | 195 | 41.7 | 27 |
| GO | dr | Bio | 96 | 10 | 35 | 14.3 | 13 |
| GO | hs | Bio | 3628 | 10 | 195 | 42.8 | 26 |
| GO | mm | Bio | 2794 | 10 | 145 | 33.0 | 23 |
| GO | sc | Bio | 1547 | 10 | 198 | 41.6 | 25 |
| MO | ce | Bio | 580 | 10 | 147 | 31.3 | 22 |
| MO | dm | Bio | 113 | 10 | 185 | 62.7 | 42 |
| MO | dr | Bio | 17 | 10 | 19 | 13.0 | 12 |
| MO | hs | Bio | 3112 | 10 | 196 | 38.2 | 23 |
| MO | mm | Bio | 2733 | 10 | 141 | 29.3 | 21 |
| MO | sc | Bio | 203 | 10 | 196 | 49.5 | 31 |
| GO | ce | IMP | 1289 | 10 | 198 | 39.5 | 23 |
| GO | dm | IMP | 2105 | 10 | 199 | 42.9 | 27 |
| GO | dr | IMP | 1205 | 10 | 196 | 40.4 | 24 |
| GO | hs | IMP | 3715 | 10 | 200 | 44.2 | 26 |
| GO | mm | IMP | 3755 | 10 | 200 | 42.3 | 26 |
| GO | sc | IMP | 1547 | 10 | 198 | 41.6 | 25 |
| MO | ce | IMP | 767 | 10 | 189 | 38.7 | 25 |
| MO | dm | IMP | 116 | 10 | 199 | 73.4 | 55 |
| MO | dr | IMP | 785 | 10 | 187 | 26.3 | 17 |
| MO | hs | IMP | 3236 | 10 | 199 | 39.0 | 24 |
| MO | mm | IMP | 3835 | 10 | 200 | 38.6 | 24 |
| MO | sc | IMP | 203 | 10 | 196 | 49.5 | 31 |

#### Optimal choices for gene classification using multi-species networks

To determine the optimal choice of network combination, representation, and hyperparameter setting that could improve the GenePlexus algorithm, we performed extensive evaluations on the tasks of predicting human and mouse annotations.

##### Which species to include?

The feature spaces were grouped into three main categories:

- **Single**: Features were only derived from network of the species the prediction task is in
- **Hs-mm**: Features were generated from jointly modeling human and mouse networks
- **All-species**: Features were generated from jointly modeling all species considered in this work (human, mouse, zebrafish, fly, worm, and yeast)

##### How to connect genes across species?

For Hs-mm and All-species, we weighted the edges for the cross-species connections based on two strategies (described in detail above):

- **Uniform**: All edge weights were set to the same value.
  - We evaluated weights equal to 1, 2, and 5.
- **Degree Weighted**: Directed edges were generated based on the within-species weighted-degree of the source node, the number of cross-species connections of the source node, and a scaling factor (see eqn. SM1).
  - The scaling parameter was set to 1.0 or 0.50.

##### How to represent the network as features in the ML model?

Within each of the three main groups listed above, the features are generated using four different methods:

- **AdjMat**: There are no additional hyperparameters.
- **RWR**: We fixed the restart parameter to 0.8 and tested using either the influence matrix directly or its transpose.
- **SVD**: We tested two settings, one with and the other without standardizing the adjacency matrix column-wise before performing SVD.
- **N2V**: Involved parameters include $p$ and $q$, which control how random walks are generated. We also performed some minimal tuning on the number of nodes per walk (*i.e.*, walk length), the number of walks per gene, and the number of epochs used when training the model.

For both SVD and N2V, we considered embedding dimension sizes of 500, 2000, and 10000. We note that, for the All-Species category, only the *node2vec* method could handle jointly modeling information from more than two species.

First, we observed that ‘degree-weighted’ connections between cross-species gene pairs (compared to ‘uniform’) resulted in more cohesion between genes in different species networks

(**Fig. 6**; **Fig. SM19–21**). So, fixing those connections, we ran all combinations of network representation methods (and their parameter sets) and species choices for function and phenotype prediction tasks. **Figure SM3** shows the results for the best hyperparameter from this analysis. This analysis required substantial computational resources and generated over 3.5TB worth of files just for human and mouse. Therefore, to compare across species other than just human and mouse, we picked one non-embedding method — AdjMat (over RWR) — and one embedding method — N2V with a $p$ = 0.1, $q$ = 0.001 and a dimension size of 2000 (over SVD) [**Fig. SM4**] — and used these two in all subsequent analyses (N2V Selected).

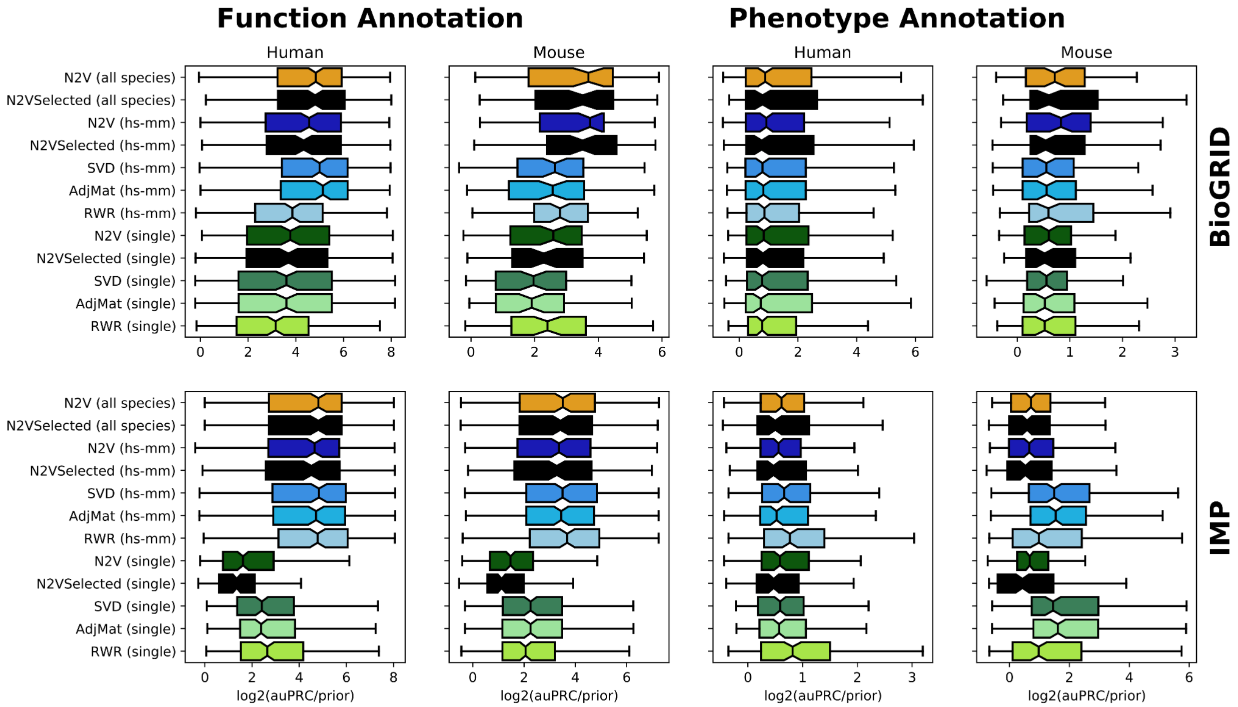


**Figure SM3: Evaluation to determine optimal choices of settings and hyperparameters for improving GenePlexus with multi-species networks**. Each group of boxplots shows the function and phenotype prediction performance for human and mouse using different species choice (for the network) and network representation methods (each using the best hyperparameter set, N2VSelected) based on networks from BioGRID and IMP.

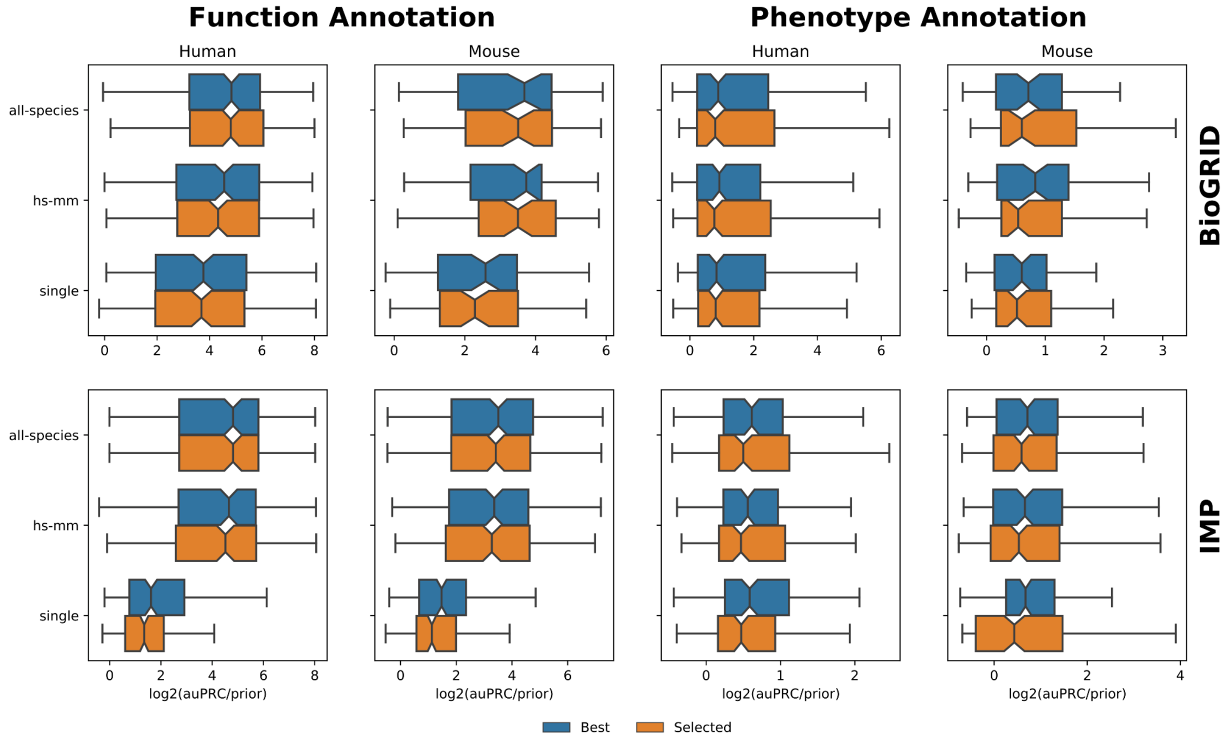


**Figure SM4: Performance of the selected hyperparameter set for node embeddings**. The boxplots show the log2(auPRC/prior) performance metric of the hyperparameter set we selected for N2V (orange) to use in the whole study compared to the performance of the best hyperparameter set (blue) across function and phenotype prediction tasks in human and mouse using networks from BioGRID and IMP.

### Adding orthologs as positives

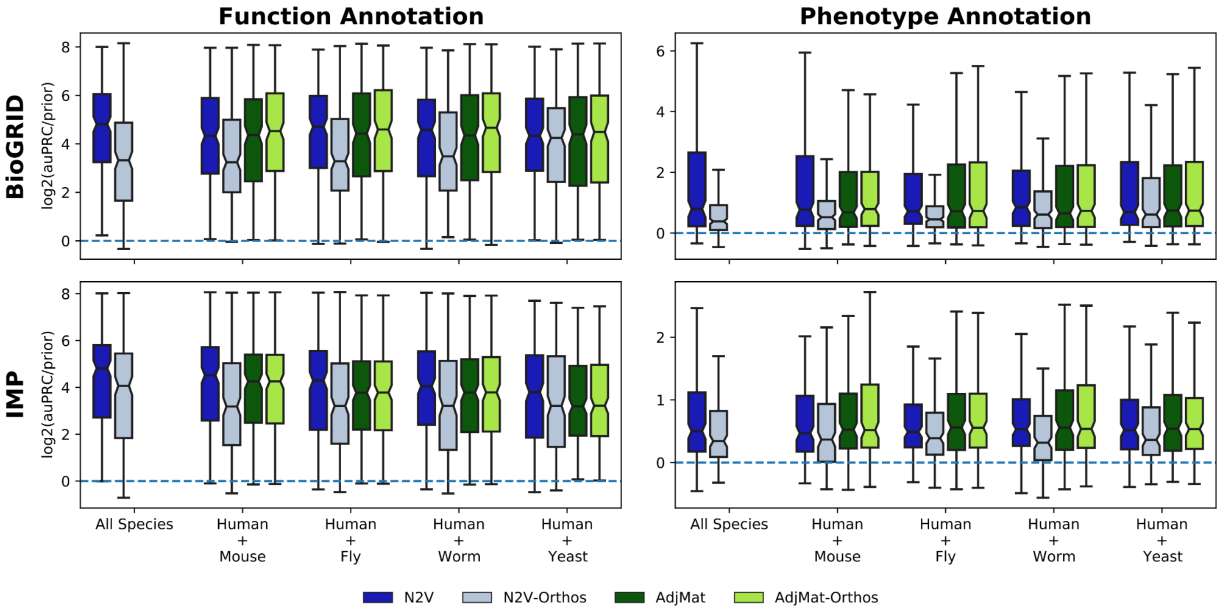


**Figure SM5: Adding orthologs for predicting function- and phenotype-associated human genes.** The same analysis as Fig. 2, but now including orthologs from the model species as additional positive examples during training.

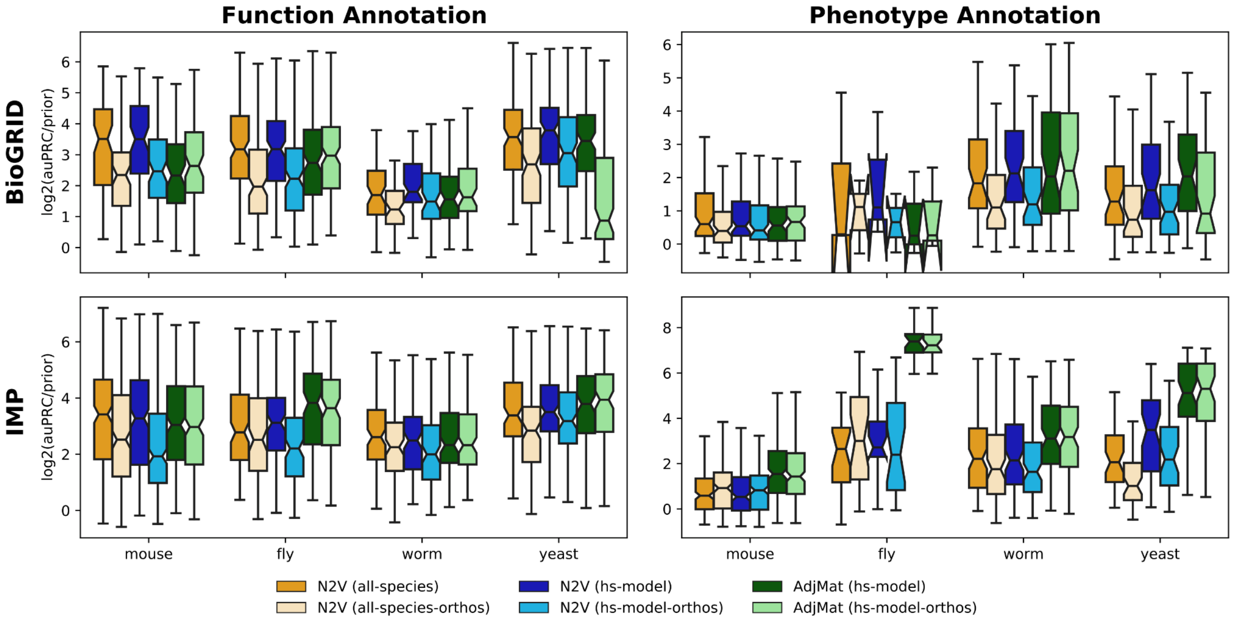


**Figure SM6: Adding orthologs for predicting function- and phenotype-associated model species genes.** The same analysis as Fig. 3, but now including orthologs from the model species as additional positive examples during training.

### Cross-species knowledge transfer

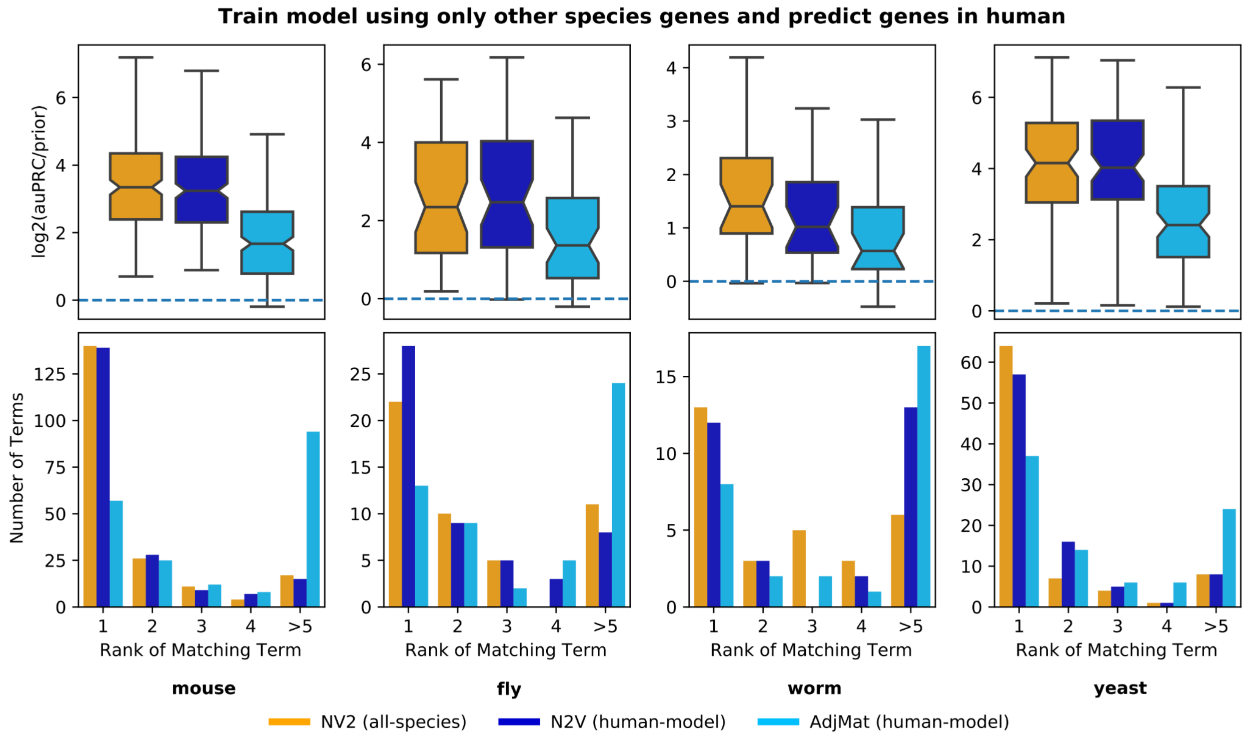


**Figure SM7: Transferring knowledge from model species to human using human+model and all-species representations of networks from IMP.** The boxplots (top row) show the $\log_2$(auPRC/prior) prediction performance of logistic regression classifiers that were trained using only model species genes annotated to a given biological process and then used to predict human genes annotated to the same biological process. The barplots (bottom row) show the counts of where the $\log_2$(auPRC/prior) for the matched biological process ranks in comparison to the other processes in the collection. Features for the classifier were created pairwise between human and a model species with IMP networks using the AdjMat (light blue) and N2V (dark blue) representations, as well as features from all species using the N2V (orange) representation.

The following two figures are the corresponding figures to Figures 4 and SM7, respectively, except generated using the BioGRID network.

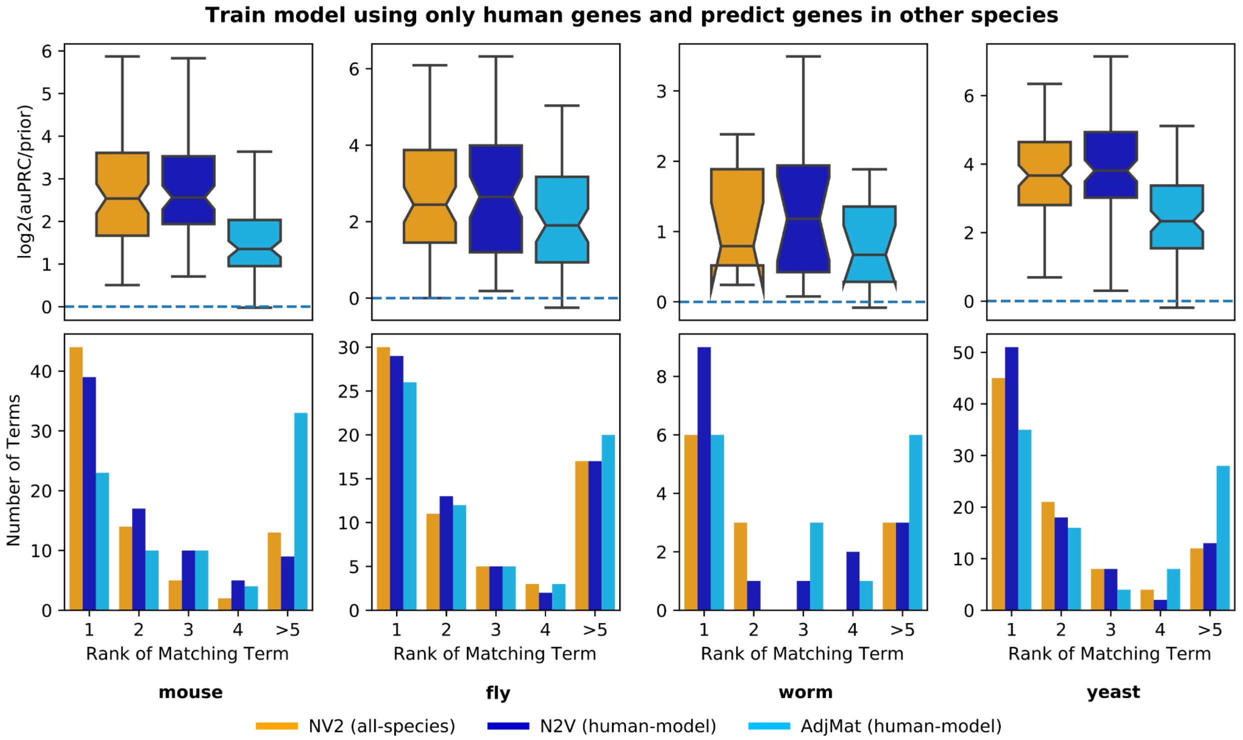


**Figure SM8: Transferring knowledge from human to model species using human+model and all-species representations of networks from BioGRID.** The boxplots (top row) show the $\log_2$(auPRC/prior) prediction performance of logistic regression classifiers that were trained using only human genes annotated to a given biological process and then used to predict model species genes annotated to the same biological process. The barplots (bottom row) show the counts of where the $\log_2$(auPRC/prior) for the matched biological process ranks in comparison to the other processes in the collection. Features for the classifier were created pairwise between human and a model species with BioGRID networks using the AdjMat (light blue) and N2V (dark blue) representations, as well as features from all species using the N2V (orange) representation.

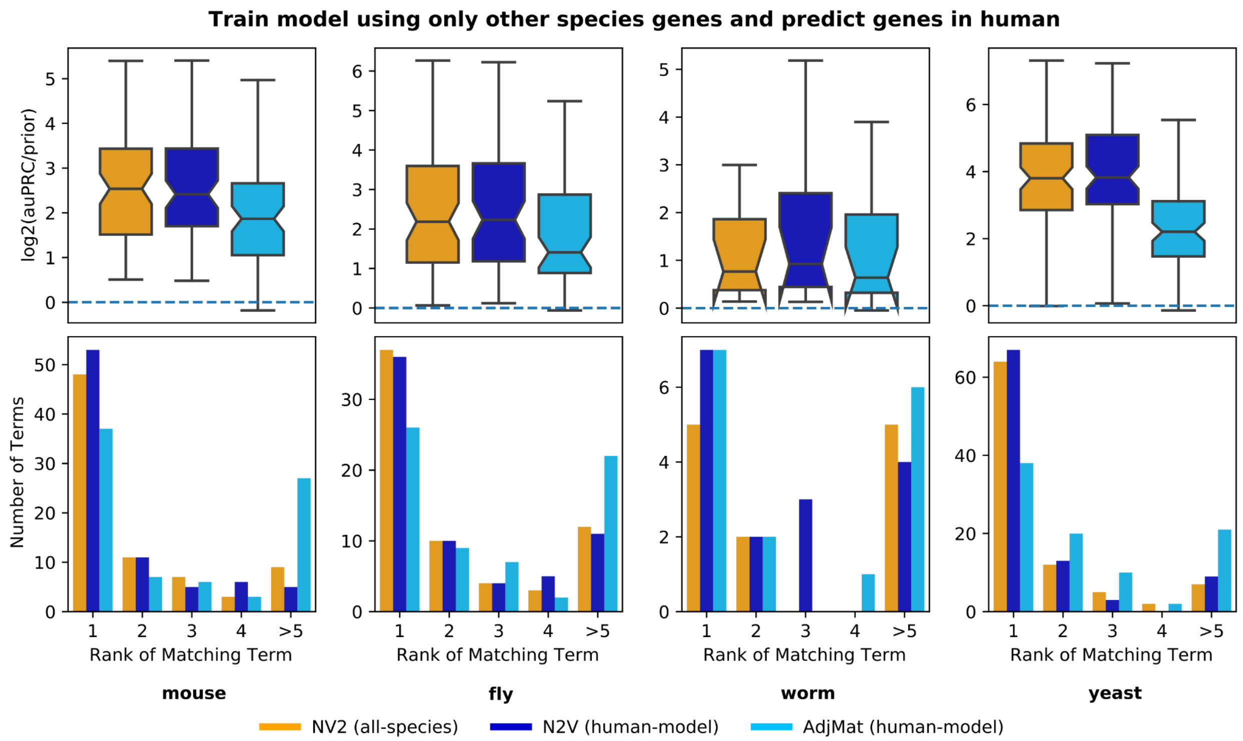


**Figure SM9: Transferring knowledge from model species to human using human+model and all-species representations of networks from BioGRID.** The boxplots (top row) show the $\log_2$(auPRC/prior) prediction performance of logistic regression classifiers that were trained using only model species genes annotated to a given biological process and then used to predict human genes annotated to the same biological process. The barplots (bottom row) show the counts of where the $\log_2$(auPRC/prior) for the matched biological process ranks in comparison to the other processes in the collection. Features for the classifier were created pairwise between human and a model species with BioGRID networks using the AdjMat (light blue) and N2V (dark blue) representations, as well as features from all species using the N2V (orange) representation.

### Same-species prediction vs. full cross-species prediction

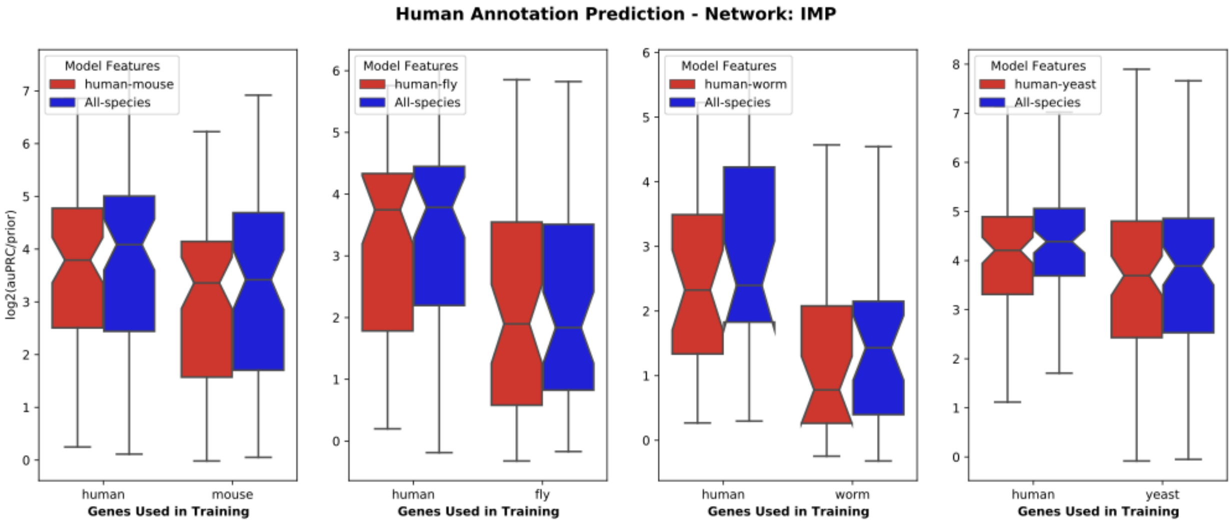


**Figure SM10: Predicting human gene annotations by training classifiers with genes only from human or only from another species using IMP networks.** Each panel corresponds to the analysis of human and one other model species. Within each panel, the boxplots show the performance on the task of predicting human genes annotated to biological processes using classifiers that were trained either on human genes only (left pair) or model species genes only (right pair), based on features from the human-model (red) or all six species (blue) networks from IMP.

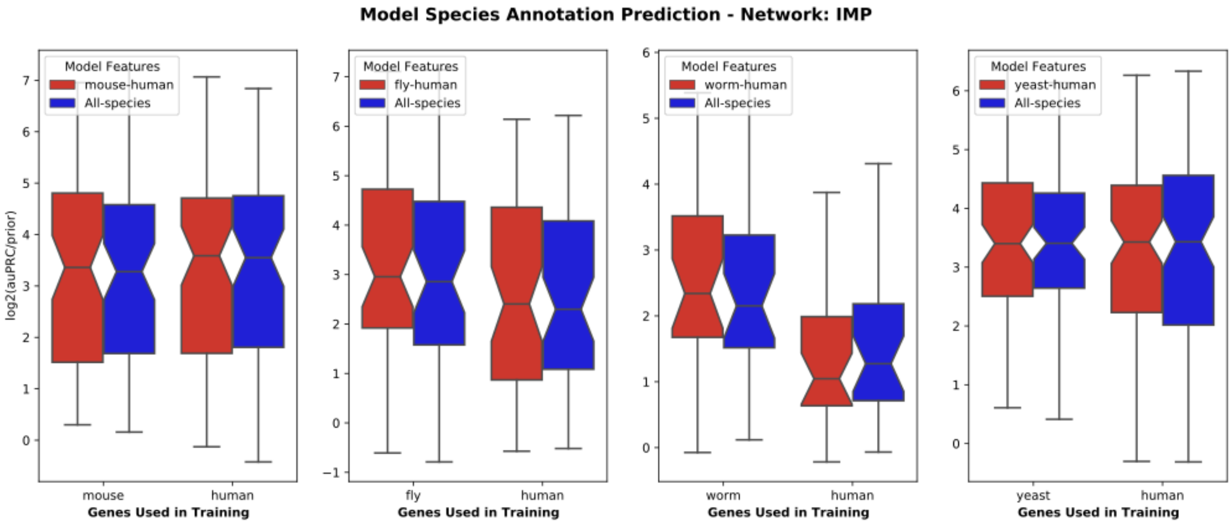


**Figure SM11: Predicting model species gene annotations by training classifiers with genes only from that species or only from human using IMP networks.** Each panel corresponds to the analysis of one model species with human. Within each panel, the boxplots show the performance on the task of predicting model species genes annotated to biological processes using classifiers that were trained either on model species genes only (left pair) or human genes only (right pair), based on features from the human-model (red) or all six species (blue) networks from IMP.

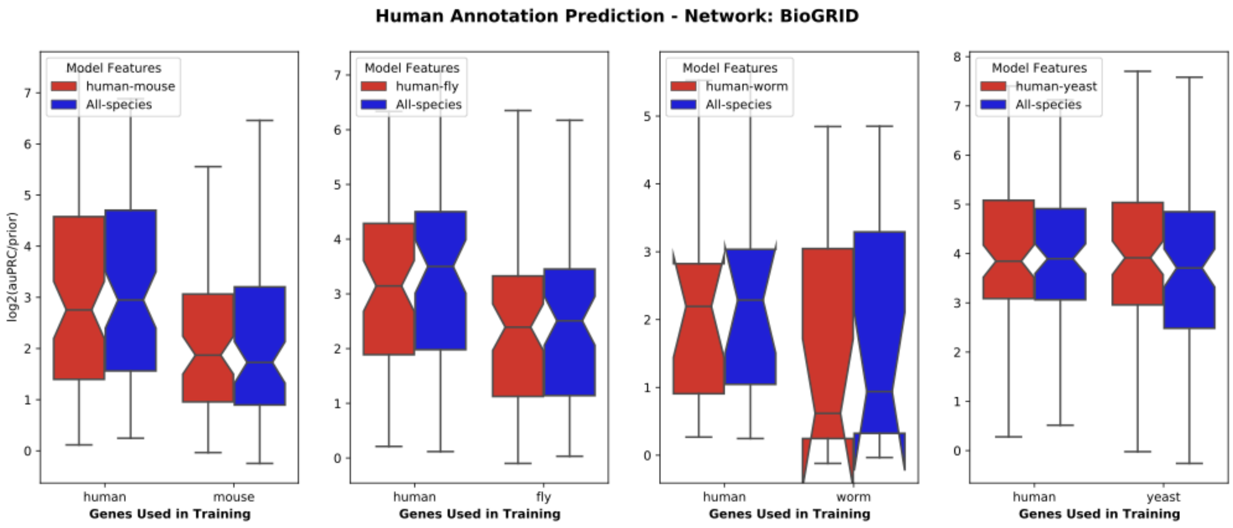


**Figure SM12: Predicting human gene annotations by training classifiers with genes only from human or only from another species using BioGRID networks.** Each panel corresponds to the analysis of human and one other model species. Within each panel, the boxplots show the performance on the task of predicting human genes annotated to biological processes using classifiers that were trained either on human genes only (left pair) or model species genes only (right pair), based on features from the human-model (red) or all six species (blue) networks from BioGRID.

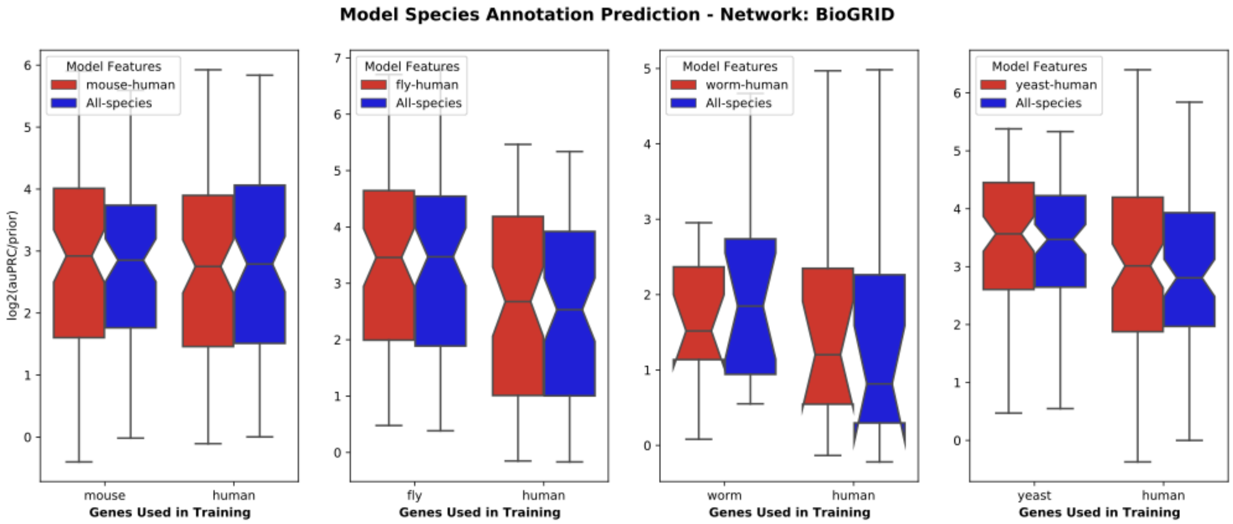


**Figure SM13: Predicting model species gene annotations by training classifiers with genes only from that species or only from human using BioGRID networks.** Each panel corresponds to the analysis of one model species with human. Within each panel, the boxplots show the performance on the task of predicting model species genes annotated to biological processes using classifiers that were trained either on model species genes only (left pair) or human genes only (right pair), based on features from the human-model (red) or all six species (blue) networks from BioGRID.

### Adding genes from multiple species during training

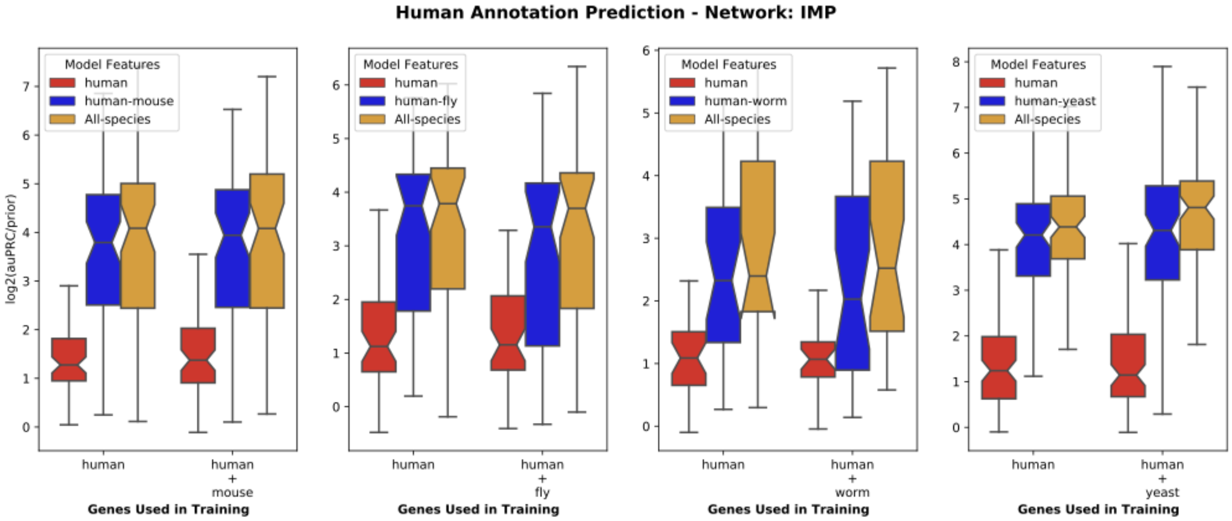


**Figure SM14: Predicting human gene annotations by including genes from multiple species during training with IMP networks.** Each panel corresponds to the analysis of human and one model species. Within each panel, the boxplots show the performance on the task of predicting human genes annotated to biological processes using classifiers that were trained either on human genes only (left pair) or human+model genes (right pair), based on features from the human (red), human+model (blue), or all six species (orange) networks from IMP.

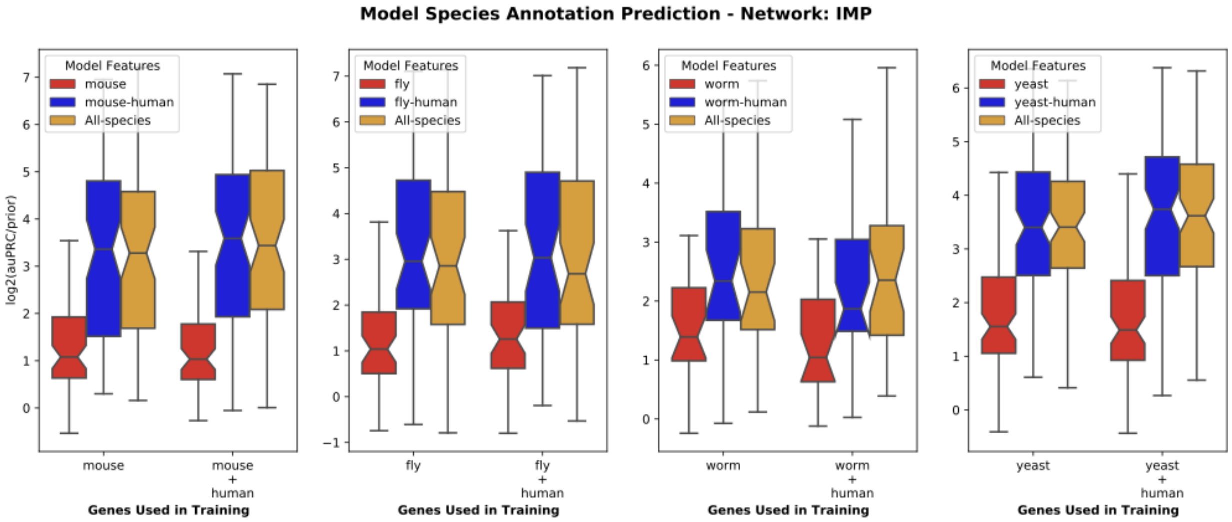


**Figure SM15: Predicting model species gene annotations by including genes from multiple species during training with IMP networks.** Each panel corresponds to the analysis of one model species with human. Within each panel, the boxplots show the performance on the task of predicting model species genes annotated to biological processes using classifiers that were trained either on model species genes only (left pair) or model+human genes (right pair), based on features from the model (red), model+human (blue), or all six species (orange) networks from IMP.

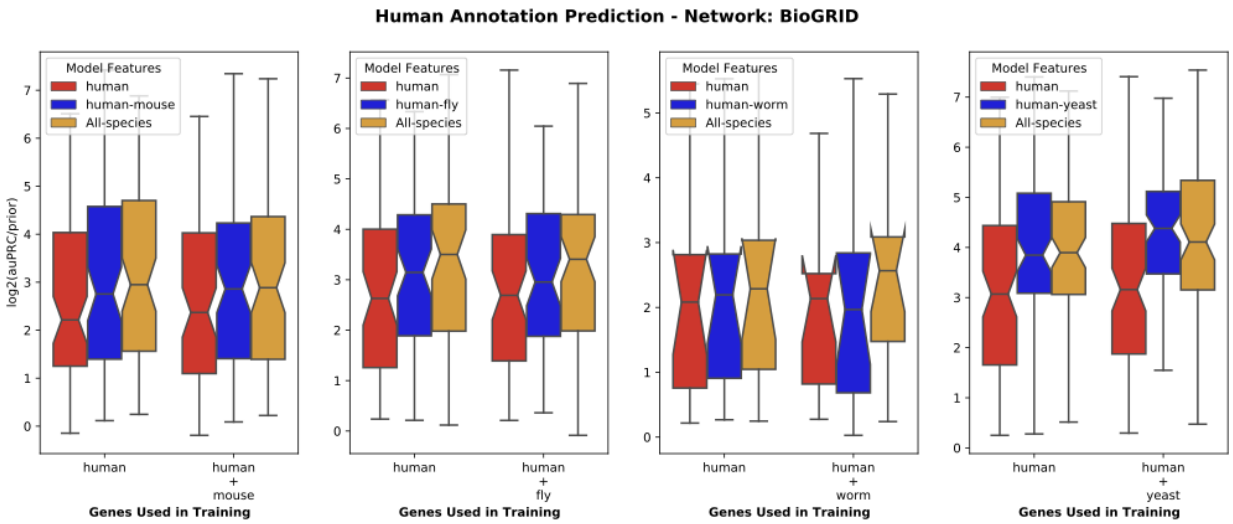


**Figure SM16: Predicting human gene annotations by including genes from multiple species during training with BioGRID networks.** Each panel corresponds to the analysis of human and one model species. Within each panel, the boxplots show the performance on the task of predicting human genes annotated to biological processes using classifiers that were trained either on human genes only (left pair) or human+model genes (right pair), based on features from the human (red), human+model (blue), or all six species (orange) networks from BioGRID.

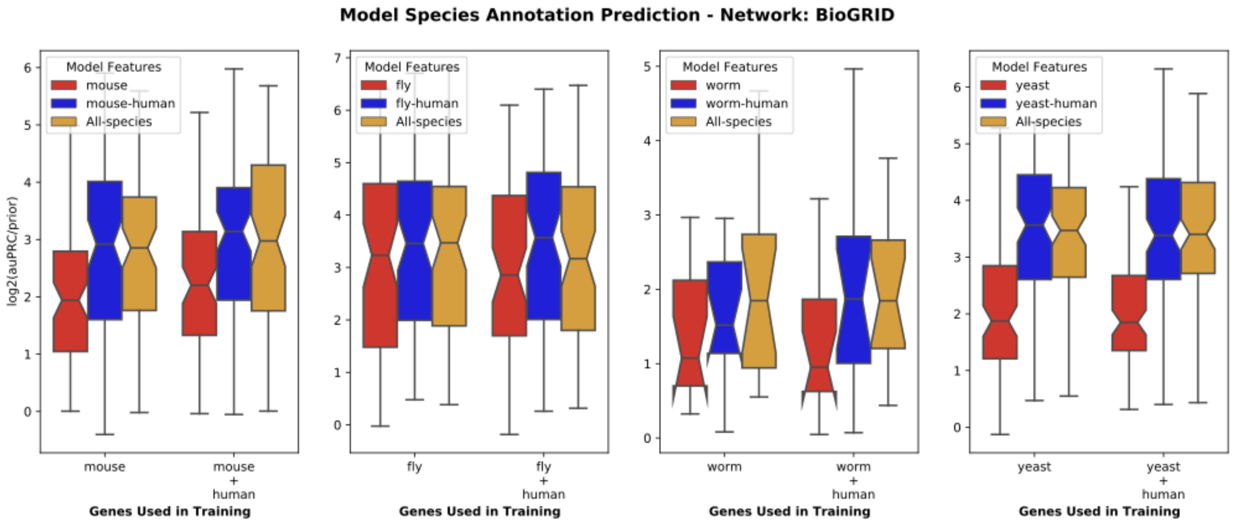


**Figure SM17: Predicting model species gene annotations by including genes from multiple species during training with BioGRID networks.** Each panel corresponds to the analysis of one model species with human. Within each panel, the boxplots show the performance on the task of predicting model species genes annotated to biological processes using classifiers that were trained either on model species genes only (left pair) or model+human genes (right pair), based on features from the model (red), model+human (blue), or all six species (orange) networks from BioGRID.

#### Predicting model species counterparts of human diseases

For a given species, the model trained on human disease gene annotations can be used to predict a probability of how associated every gene in all six species is to the human disease. To determine the z-score for the *i*-th gene, we use:

$$z_{gene,i} = p_i - \mu_p / \sigma_p \ , \qquad \text{(eqn. SM3)}$$

where $p_i$ is the probability of the *i*-th gene, $\mu_p$ and $\sigma_p$ are the mean value and standard deviation of the probability distribution for a given species, respectively. To calculate the z-score for a GO or Monarch gene set, we use:

$$z_{set} = (\mu_{set} - \mu_p) / (\sigma_p / \sqrt{n}) \ , \qquad \text{(eqn. SM4)}$$

where $\mu_{set}$ is the mean value of probabilities for only genes annotated to the set, $n$ is the number of genes in the set, and $\mu_p$ and $\sigma_p$ are the mean value and standard deviation of the probability distribution for a given species, respectively.

**Table SM5: Bardet-Biedl Syndrome 1 top ten predicted genes in each species.** Top ten genes in each species (columns) based on the z-score of the prediction probabilities from the model. Genes in each species are ranked independently, *i.e.*, genes from different species in the same row are not intended to correspond to each other.

| Bardet-Biedl syndrome top ten associated genes | | | | | |
|---|---|---|---|---|---|
| human | mouse | fish | fly | worm | yeast |
| Bardet-Biedl syndrome 9 | Bardet-Biedl syndrome 2 (human) | Bardet-Biedl syndrome 5 | Meckel syndrome, type 1 | Bardet-Biedl syndrome 7 protein homolog | chaperonin-containing T-complex subunit CCT7 |
| Bardet-Biedl syndrome 7 | Bardet-Biedl syndrome 5 (human) | nephronophthisis 1 | Bardet-Biedl syndrome 5 | Tetratricopeptide repeat protein 8 | chaperonin-containing T-complex subunit CCT5 |
| Bardet-Biedl syndrome 2 | tetratricopeptide repeat domain 8 | tetratricopeptide repeat domain 21B | Outer segment 2 | Bardet-Biedl syndrome 2 protein homolog | 1-phosphatidylinositol-3-phosphate 5-kinase |
| Bardet-Biedl syndrome 4 | Bardet-Biedl syndrome 9 (human) | Bardet-Biedl syndrome 7 | Bardet-Biedl syndrome 9 | Protein pthb1 homolog | HSP70/90 family co-chaperone CNS1 |
| tetratricopeptide repeat domain 8 | centrosomal protein 290 | intraflagellar transport 172 | Bardet-Biedl syndrome 8 | Nephrocystin-1-like protein | chaperonin-containing T-complex alpha subunit TCP1 |
| Bardet-Biedl syndrome 5 | intraflagellar transport 172 | Bardet-Biedl syndrome 2 | Bardet-Biedl syndrome 1 | MecKel-Gruber Syndrome (MKS) homolog | Kar3p |
| tetratricopeptide repeat domain 21B | Bardet-Biedl syndrome 1 (human) | MKS transition zone complex subunit 1 | Heat shock protein 60C | Bardet-Biedl syndrome 5 protein homolog | chaperonin-containing T-complex subunit CCT3 |
| centrosomal protein 290 | tetratricopeptide repeat domain 21B | centrosomal protein 290 | Heat shock protein 60B | MecKel-Gruber Syndrome (MKS) homolog | chaperonin-containing T-complex subunit CCT2 |
| Bardet-Biedl syndrome 1 | MKS transition zone complex subunit 1 | Bardet-Biedl syndrome 1 | uncharacterized protein | Intraflagellar transport protein osm-1 | lysophospholipase |
| leucine zipper transcription factor like 1 | Bardet-Biedl syndrome 7 (human) | dynein, cytoplasmic 2, light intermediate chain 1 | Intraflagellar transport 46 | X-BoX promoter element regulated | transcription regulator CYC8 |

**Table SM6: Bardet-Biedl Syndrome 1 top ten associated biological processes in each species.** Top ten associated GO biological processes in each species (columns) based on the z-score of the predicted probability of the genes in each species. Processes in each species are ranked independently, *i.e.*, processes from different species in the same row are not intended to correspond to each other. Real relationships between these processes are presented in Figure 5.

| Bardet-Biedl syndrome top ten associated biological processes | | | | | |
|---|---|---|---|---|---|
| human | mouse | fish | fly | worm | yeast |
| protein localization to cilium | cilium organization | melanosome transport | non-motile cilium assembly | cilium organization | protein folding |
| intraciliary transport involved in cilium assembly | non-motile cilium assembly | establishment of melanosome localization | cilium assembly | intraciliary transport | spindle assembly |
| intraciliary transport | cilium assembly | pigment granule transport | cilium organization | protein transport along microtubule | protein depolymerization |
| microtubule-based protein transport | regulation of cilium-dependent cell motility | melanosome localization | plasma membrane bounded cell projection assembly | microtubule-based protein transport | protein refolding |
| protein transport along microtubule | retina homeostasis | intracellular transport | cell projection assembly | cilium assembly | mitotic nuclear division |
| transport along microtubule | intraciliary transport | establishment of pigment granule localization | regulation of establishment of planar polarity | plasma membrane bounded cell projection assembly | mitotic spindle elongation |
| microtubule-based transport | striatum development | establishment of vesicle localization | axoneme assembly | cell projection assembly | spindle elongation |
| cytoskeleton-dependent intracellular transport | photoreceptor cell maintenance | pigment granule localization | microtubule bundle formation | protein-containing complex localization | glycerolipid metabolic process |
| microtubule-based movement | smoothened signaling pathway | vesicle localization | regulation of morphogenesis of an epithelium | microtubule-based transport | organelle transport along microtubule |
| ciliary basal body-plasma membrane docking | protein transport along microtubule | establishment of localization in cell | microtubule-based movement | transport along microtubule | transport along microtubule |

**Table SM7: Bardet-Biedl Syndrome 1 top ten associated phenotypes in each species.** Top ten associated Monarch phenotypes in each species (columns) based on the z-score of the predicted probability of the genes in each species. Phenotypes in each species are ranked independently, *i.e.*, phenotypes from different species in the same row are not intended to correspond to each other.

| Bardet-Biedl syndrome top ten associated phenotypes | | | | | |
|---|---|---|---|---|---|
| human | mouse | fish | fly | worm | yeast |
| Medial flaring of the eyebrow | photoreceptor inner segment degeneration | gastrulation disrupted, abnormal | abnormal planar polarity | ivermectin resistant | spindle morphology:abnormal |
| Hypoplasia of the ovary | absent sperm flagellum | notochord kinked, abnormal | lethal - all die during pharate adult stage | osmotic avoidance defective | chromosome segregation:abnormal |
| Postaxial hand polydactyly | Retinal degeneration | pronephric proximal convoluted tubule morphology, abnormal | sensory perception of sound process quality, abnormal | abnormal cilium morphology | nuclear position:abnormal |
| Hepatic fibrosis | retinal outer nuclear layer degeneration | pronephros development disrupted, abnormal | male semi-sterile | associative learning variant | vacuolar transport:decreased |
| Generalized hirsutism | enlarged third ventricle | somite increased width, abnormal | majority die during pharate adult stage | dye filling defect | vacuolar transport:abnormal |
| Multicystic kidney dysplasia | absent embryonic cilia | notochord increased width, abnormal | some die during first instar larval stage | vulva location variant | chemical compound accumulation:absent |
| Nephrotic syndrome | disorganized photoreceptor outer segment | convergent extension process quality, abnormal | cytokinesis process quality, abnormal | male response to contact defective | protein secretion:decreased |
| Polydactyly | abnormal renal tubule epithelial cell primary cilium morphology | pronephros cystic, abnormal | semi-fertile | ammonium chloride chemotaxis defective | cytoskeleton morphology:abnormal |
| Pigmentary retinopathy | small hippocampus | pronephric duct dilated, abnormal | sensory perception of touch process quality, abnormal | cephalic sensillum morphology variant | freeze-thaw resistance:decreased |
| External genital hypoplasia | abnormal motile primary cilium morphology | cilium Kupffer's vesicle decreased length, abnormal | photoperiod response variant | male mating efficiency reduced | protein activity:absent |

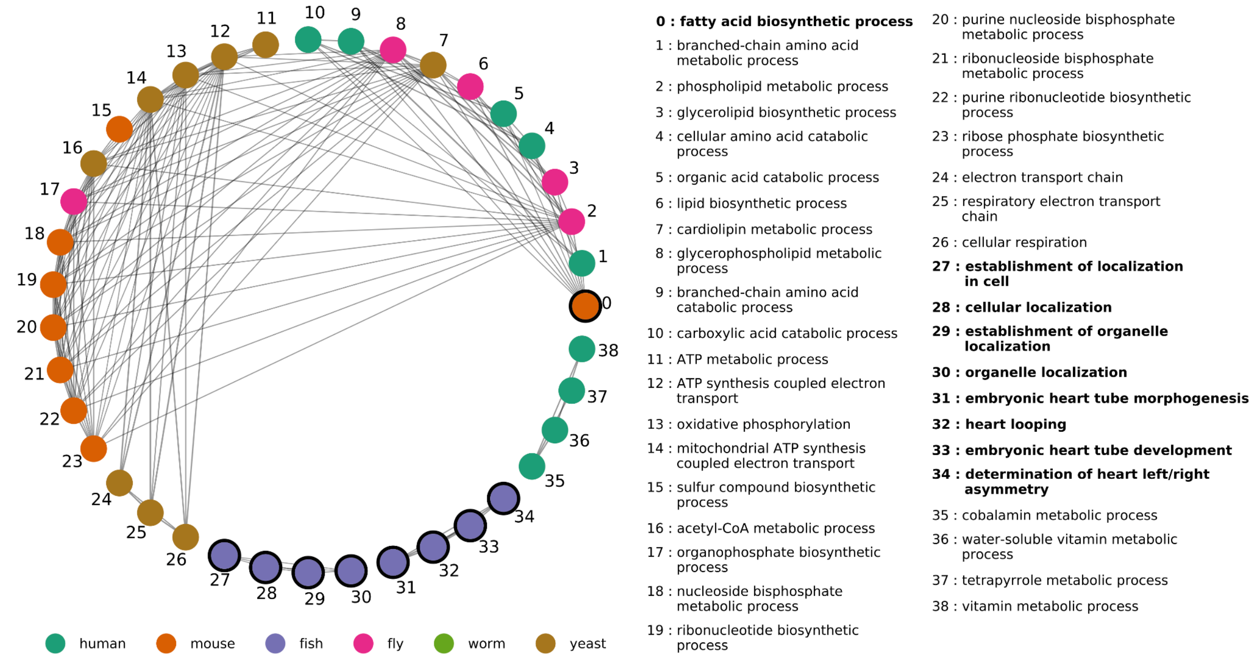


**Figure SM18: The ten most enriched biological processes associated with the top genes in each species predicted to be related to organic acidemia (OA).** A classifier trained using human OA genes and all-species N2V features was used to predict OA-related genes in the five model species. The figure shows the ten most enriched biological processes associated with the top-ranked genes in each species. Nodes represent biological processes and are colored by which species they were identified in. Edges represent pairs of semantically similar processes (scaled Resnik similarity based on the Gene Ontology). Isolated nodes are not shown. Nodes with thick borders represent biological processes where, in at least one species, none of the annotated genes are orthologous to any human OA gene.

### Additional PCA plots

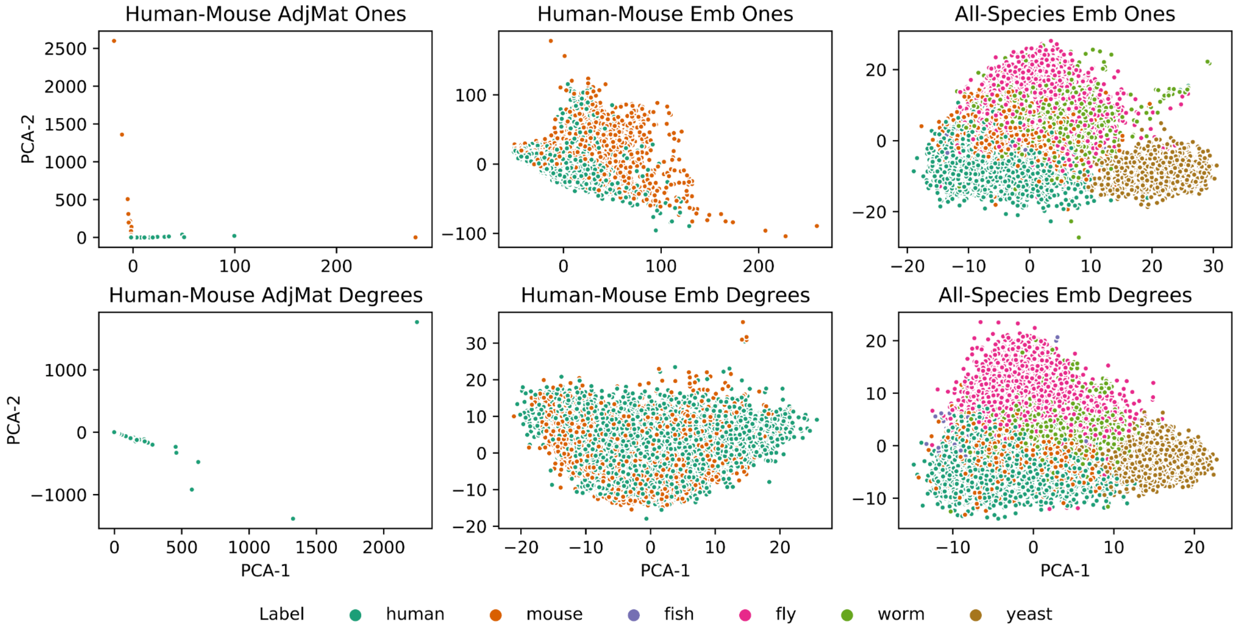


**Figure SM19: PCA of adjacency matrix and node embedding representations of human-mouse and all-species networks.** Each PCA plot shows the top two PCA components from the human-mouse adjacency matrix (first column), the human-mouse node embedding (second column) and the all species node embedding (third column) representations. The cross-species edge weights were created by always using a uniform edge weight with a value of one (top row) or by using degree based edge weights (bottom row) for the BioGRID network.

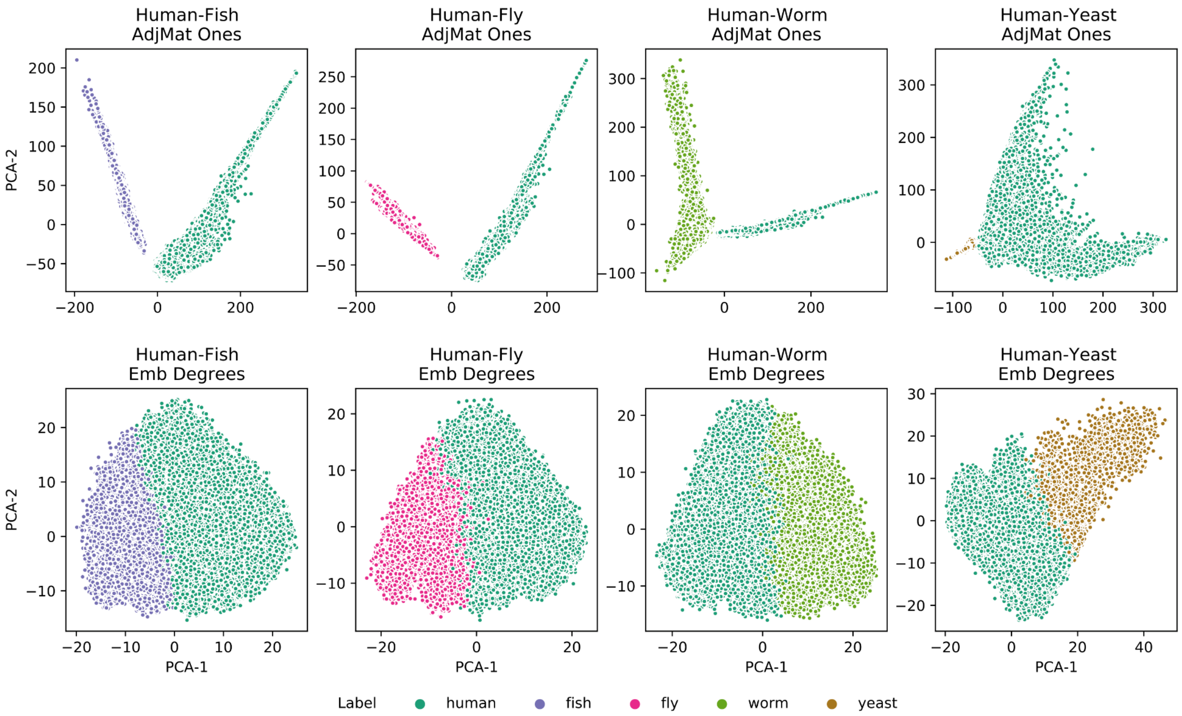


**Figure SM20: PCA of adjacency matrix and node embedding representations of human-model networks from IMP.** Each PCA plot shows the top two PCA components from the adjacency matrix (using the uniform weighting strategy; top row) and node embeddings (using the degree based weighting strategy; bottom row) representations for the human network combined pairwise with fish, fly, worm, yeast in IMP.

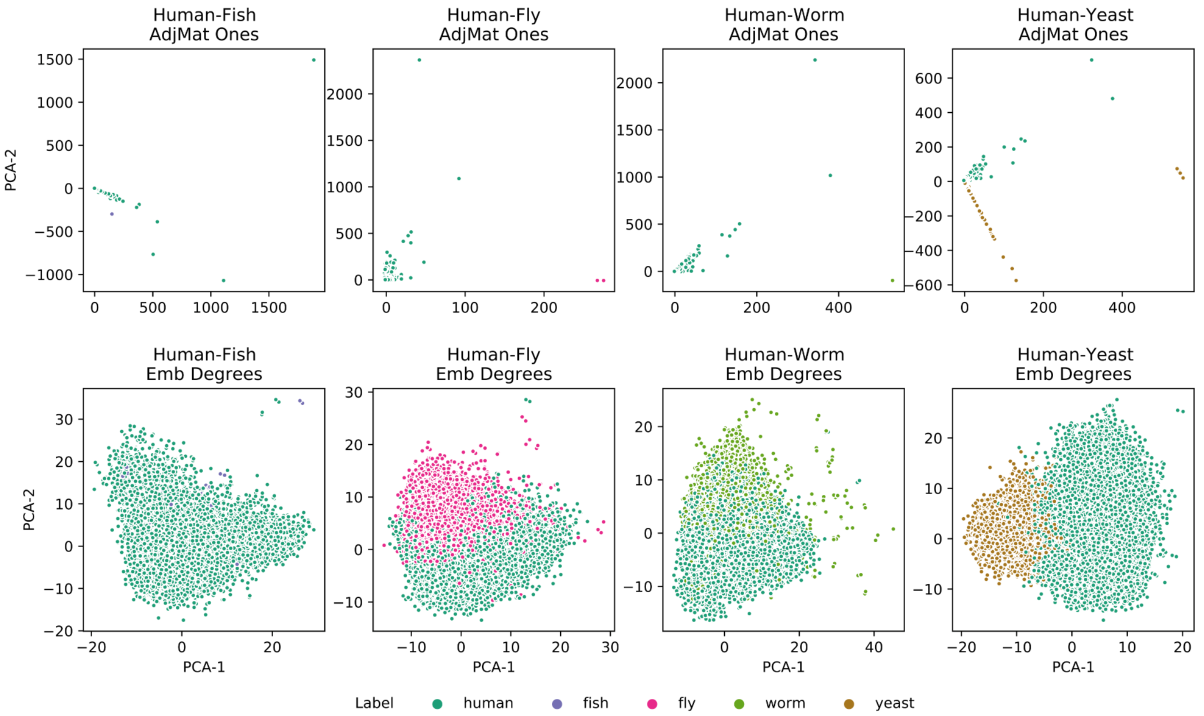


**Figure SM21: PCA of adjacency matrix and node embedding representations of human-model networks from BioGRID.** Each PCA plot shows the top two PCA components from the adjacency matrix (using the uniform weighting strategy; top row) and node embeddings (using the degree based weighting strategy; bottom row) representations for the human network combined pairwise with fish, fly, worm, yeast in BioGRID.
